# Distinct functions of PAXX and MRI during chromosomal end joining

**DOI:** 10.1101/2024.08.21.607864

**Authors:** Metztli Cisneros-Aguirre, Felicia Wednesday Lopezcolorado, Xiaoli Ping, Ruby Chen, Jeremy M. Stark

## Abstract

A key step of Canonical Nonhomologous End Joining (C-NHEJ) is synapsis of DNA double strand break (DSB) ends for ligation. The DNA-PKcs dimer mediates synapsis in a long-range complex with DSB ends remaining apart, whereas the XLF homodimer can mediate synapsis in both long-range and short-range complexes. Recent structural studies found the PAXX homodimer may also facilitate synapsis in long-range complexes with DNA-PKcs via its interactions with Ku70. Thus, we examined the influence of PAXX in C-NHEJ of chromosomal DSBs, which we compared to another Ku-binding factor, MRI. Using EJ of blunt DSBs with Cas9 reporters as a readout for C-NHEJ, we found that PAXX and/or MRI are dispensable. However, when combined with disruption of DNA-PKcs, particularly with DNA-PKcs kinase inhibition, PAXX becomes important for blunt DSB EJ. In contrast, while DNA-PKcs is also important to suppress short deletion mutations with microhomology, this effect is not magnified with PAXX loss. MRI loss had no effect combined with DNA-PKcs disruption, but becomes important for blunt DSB EJ when combined with disruption of XLF, as is PAXX. Finally, XLF loss causes an increase in larger deletions compared to DNA-PKcs inhibition, which is magnified with combined loss of MRI. Altogether, we suggest that PAXX promotes DSB end synapsis during C-NHEJ in a manner that is partially redundant with DNA-PKcs and XLF, whereas MRI appears to be mainly important in the context of XLF disruption.

**SIGNIFICANCE STATEMENT:** Canonical Nonhomologous End Joining (C-NHEJ) is a DNA double strand break repair pathway that can repair diverse types of DNA ends, but is prone to causing genetic mutations. In particular, C-NHEJ can cause insertion or deletion mutations that can cause human disease. Therefore, it is critical to understand the mechanisms of C-NHEJ, including and how various factors within the pathway affect repair outcomes. In this study, we elucidate the role of factors PAXX and MRI during chromosomal EJ and their interplay with the C-NHEJ factors DNA-PKcs and XLF.

## INTRODUCTION

Chromosomal DNA double strand breaks (DSBs) can be caused by topoisomerases, oxidative stress, various forms of replication stress, environmental clastogens, and several cancer therapeutics (1–3). End joining (EJ) is one of the major mechanisms for DNA DSB repair and is commonly associated with insertions/deletions (indels) or deletion rearrangement mutations (4–8). Indels and deletion rearrangements can cause disruption in tumor suppressor genes (9, 10) and therefore, characterizing EJ repair and mutagenesis provides insight into cancer etiology.

Canonical-nonhomologous end joining (C-NHEJ) is the major EJ DSB repair pathway and requires factors such as Ku70/80, DNA-PKcs, XRCC4, and XLF (4, 11). The C-NHEJ factors create repair complexes for the synapsis of DNA DSB ends, followed by ligation via Ligase 4 (LIG4) (12–14). C-NHEJ can repair a diverse set of DNA breaks, however other compensatory Alternative EJ (Alt-EJ) pathways such as microhomology mediated end joining (MMEJ)/theta mediated end joining (TMEJ) can also mediate such repair (4, 6, 15, 16). In the absence of C-NHEJ, Alt-EJ pathways remain proficient, but the EJ junctions show elevated deletion mutations with microhomology (4, 7, 17–20). Conversely, C-NHEJ appears required for repair of DNA DSBs that do not require an annealing intermediate, such as blunt DNA ends, or deletions involving limited (e.g., 1 nt) use of microhomology (4, 5, 20, 21). For example, repair of blunt DNA DSBs induced by Cas9 is dependent on several C-NHEJ factors, such as Ku70, XRCC4, and XLF (20, 22). Such blunt DNA EJ can be assessed by monitoring repair of two tandem Cas9 blunt DSBs causing precise incision between the DSBs, or and/or fill-in of 5’ overhangs of staggered Cas9 DSBs (a minor product of Cas9), which causes short insertions (5, 22, 23). Accordingly, these blunt DSB EJ events are robust genetic assays for C-NHEJ (5, 22, 23).

C-NHEJ appears to involve distinct Long-Range (LR) complex and Short-Range (SR) complexes, referring to the relative distances of the DSB ends. Several types of LR complexes can be visualized using Cryo-EM, where synapsis of DSB ends is mediated by the DNA-PKcs dimer (13, 14, 24). Synapsis by DNA-PKcs appears to be facilitated by the dimerization interfaces between each protomer, along with an interaction of DNA-PKcs with the Ku80 C-terminus on the opposing DSB end (13–15). Furthermore, these dimerization interfaces appear to be critical for the kinase domain of DNA-PKcs to be in close proximity to phosphorylation domains of the other DNA-PKcs protomer (14). In addition to DNA-PKcs, the XLF homodimer also appears to mediate synapsis in the LR and SR complex through interactions with XRCC4 and Ku80 (13, 14, 24). These structures suggest possible redundant synapsis mechanisms within the C-NHEJ machinery. Indeed, using Cas9 assays to measure blunt DSB EJ, DNA-PKcs was shown to play a minor role during blunt DSB EJ, however weakening XLF (i.e., disruption of binding interfaces) magnified the role for DNA-PKcs to promote such EJ events (22). These findings raise the possibility of other possible redundant interactions within the LR complex to promote chromosomal EJ.

Recent structures of the LR complex have implicated the accessory C-NHEJ factor PAXX as another possible synapsis factor (25). The PAXX homodimer appears to mediate synapsis via interaction of each protomer with Ku70 through its conserved Ku Binding Motif (KBM) (25). In addition to PAXX, MRI is another C-NHEJ accessory factor that interacts with the Ku heterodimer (26, 27), however its role in chromosomal EJ remains unclear. Both PAXX and MRI are dispensable for V(D)J recombination, but are important for this event when combined with XLF loss (27–31). However, there are two key distinctions between V(D)J recombination vs. repair of blunt Cas9 DSBs: the former requires opening of hairpin coding DNA ends, whereas the latter is dependent on XLF (7, 20). Thus, we sought to define the role of PAXX and MRI during blunt DSB EJ and elucidate possible redundancies with DNA-PKcs and XLF.

## RESULTS

### Inhibiting DNA-PKcs kinase activity reveals a role for PAXX, but not MRI, to promote No Indel EJ

We sought to define the influence of PAXX and MRI on DNA double strand break (DSB) end joining (EJ). To begin with, we examined EJ repair between two Cas9 blunt DSBs that are joined without insertion/deletion mutations from the edge of the DSB (i.e., No Indel EJ). This EJ outcome is dependent on several C-NHEJ factors (e.g., Ku70, XRCC4, and XLF), likely because C-NHEJ is uniquely capable at joining blunt DSBs (20). In contrast, theta-mediated EJ (TMEJ) requires a microhomology annealing intermediate, and hence cannot mediate No Indel EJ (20). To examine such EJ, we used the EJ7-GFP reporter chromosomally integrated into HEK293 cells (Figure 1A). This reporter contains a green fluorescent protein (GFP) sequence that is interrupted by a 46-base pair (bp) insert (7, 20, 32). Expression of Cas9 and two sgRNAs induces two DSBs that excise this insert sequence. If the two distal blunt ends are brought together for No Indel EJ, GFP is restored, leading to GFP+ cells that can be measured with flow cytometry (20).

**Figure 1.**
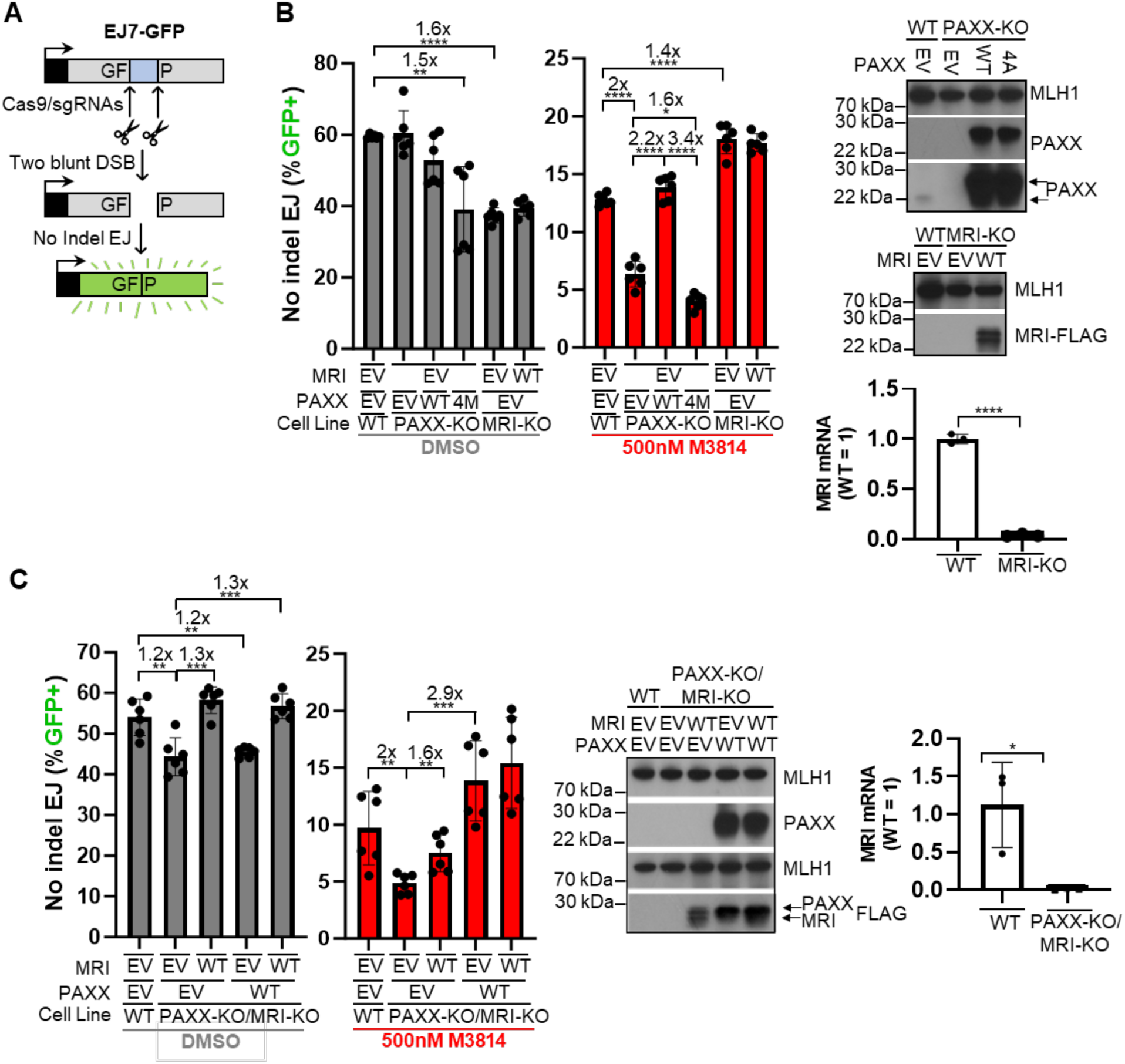
Inhibiting DNA-PKcs kinase activity reveals a role for PAXX, but not MRI, to promote No Indel EJ. **A)** Schematic of the chromosomally integrated EJ7-GFP reporter which measures No Indel EJ (i.e., blunt DSB EJ between distal ends of two Cas9 DSBs, not to scale). GFP frequencies are normalized to transfection efficiency of parallel GFP transfections. The parental cell line for the study is HEK293. **(B)** PAXX is important for No Indel EJ when combined with DNA-PKcs inhibition (M3814 treatment), whereas MRI has no obvious effect. PAXX-4M is the S184A/N187E/V199A/F201A mutant, which disrupts the Ku70 binding motif. n=6 biologically independent transfections. Statistics with unpaired t-test using Holm-Sidak correction. *P<0.05, **P<0.01, ****P<0.0001. Immunoblots show expression levels of PAXX and MRI-FLAG. RT-PCR analysis of MRI mRNA levels are shown. **(C)** Combined disruption of PAXX and MRI shows similar effects as the PAXX single disruption for No Indel EJ with M3814 treatment. n=6 biologically independent transfections. Statistics with unpaired t-test using Holm-Sidak correction. **P<0.01, ***P<0.001. Immunoblots show expression levels of PAXX, PAXX-FLAG, and MRI-FLAG. RT-PCR analysis of MRI mRNA levels is shown.

With the EJ7-GFP reporter, we examined No Indel EJ frequencies in PAXX-KO and MRI-KO cells that were treated with or without 500 nM M3814, which is a potent and specific DNA-PKcs kinase inhibitor that competes with ATP binding (33–35). Inhibition of DNA-PKcs kinase activity blocks its autophosphorylation and appears to prevent its dissociation from DNA ends (33–35). We found that PAXX-KO cells were not statistically different from WT for No Indel EJ (Figure 1B). However, comparing these two cell lines treated with 500 nM of M3814, PAXX-KO cells showed a 2-fold decrease in No Indel EJ compared to WT cells, which was rescued by transient expression of PAXX-WT (Figure 1B). Furthermore, expressing a PAXX mutant of the Ku70 binding motif (PAXX-4M: S184A/N187E/V199A/F201A) failed to restore No Indel EJ (3.4-fold difference vs. PAXX-WT). Indeed, including the 4M mutant caused a decrease compared to the empty vector control (1.5-fold), indicating an inhibitory effect of the mutant. With MRI-KO cells, we found a modest, but significant 1.6-fold decrease in No Indel EJ vs. WT (Figure 1B). In contrast to PAXX, with 500 nM M3814 treatment, MRI-KO failed to show a defect vs. WT, and indeed No Indel EJ was modestly higher than WT (1.4-fold). Notably, expression of MRI-WT in MRI-KO cells did not obviously have an effect on No Indel EJ. Finally, we considered that combined loss of PAXX and MRI might have effects on No Indel EJ, and so we generated a PAXX and MRI double KO cell line (PAXX-KO/MRI-KO). However, we found similar results with this double KO cell line as the PAXX-KO cell line (Figure 1C).

As controls, we examined levels of DNA-PKcs phosphorylation following ionizing radiation (IR) treatment, with and without 500 nM M3814, and found no effects of loss of PAXX and MRI (Supplemental Figure 1A, 1B). For the KO lines, we confirmed loss of PAXX by immunoblot, and note that PAXX complementation causes overexpression of PAXX, and confirmed loss of MRI via qRTPCR (Figure 1B, 1C). Overall, these results suggest that DNA-PKcs kinase inhibition reveals a role for PAXX, and the PAXX-KBM, to promote No Indel EJ, but not for MRI.

### DNA-PKcs kinase inhibition has a greater effect on EJ outcomes when combined with PAXX loss

We next tested the above hypothesis using an assay that can examine various EJ outcomes (Figure 2A). Specifically, we developed the MTAP/CDKN2B-AS1 deletion (MA-del) assay, which is designed to mimic deletion of the *CDKN2A* gene that is a recurrent rearrangement in many cancers, and for which the breakpoints are often in the flanking MTAP and CDKN2B-AS1 genes (10, 36). We introduced two Cas9/sgRNAs plasmids that target the MTAP and the CDK2NB-AS1 loci along with a puromycin resistance plasmid, transfected cells are enriched with puromycin selection, and the deletion rearrangement is then amplified using PCR. These PCR products are analyzed via deep sequencing, which were aligned to the predicted No Indel EJ product (i.e., joining of the distal blunt DSBs without indels, Supplemental Figure 2A). Then, the reads are categorized as No Indel, Insertions, Deletions, Complex Indels.

**Figure 2.**
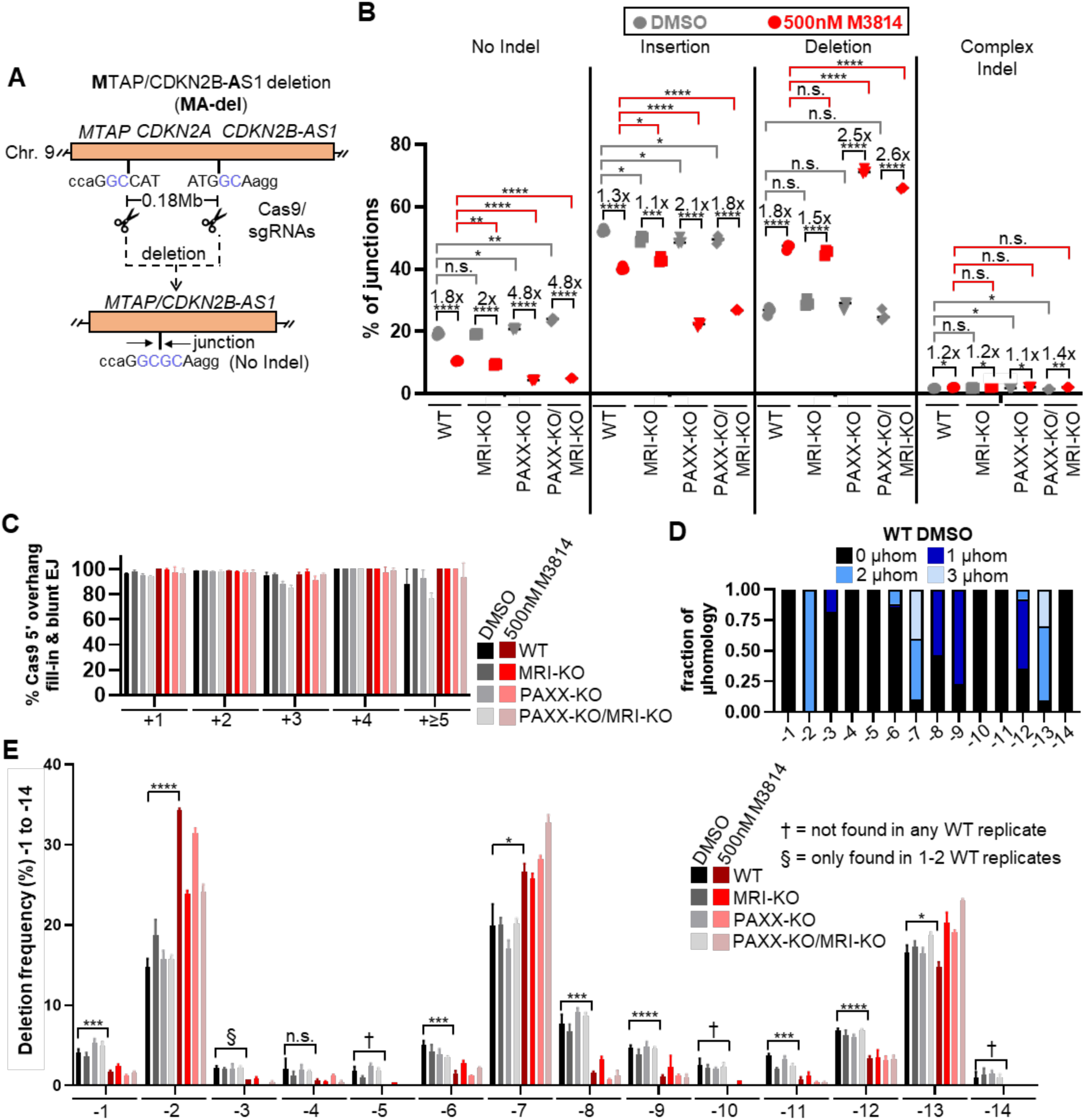
DNA-PKcs kinase inhibition has a greater effect on EJ outcomes when combined with PAXX loss. **(A)** Schematic of the MTAP/CDK2NB1-AS1 deletion rearrangement (MA-del) reporter (not to scale), which involves inducing two Cas9 DSBs at endogenous genes flanking the *CDKN2A* locus. The deletion rearrangement is amplified by PCR, which is used for deep sequencing analysis. **(B)** DNA-PKcs inhibition (M3814 treatment) causes a reduction in No Indel EJ and Insertions and a converse increase in Deletions, and these effects are greater in PAXX-KO cells. Shown are junction frequencies for No Indel EJ, Insertions, Deletions, and Complex Indels for each condition. n=3 biologically independent transfections. Statistics with unpaired t-test using Holm-Sidak correction. *P<0.05, **P<0.01, P<0.001, ****P<0.0001, n.s. = not significant. **(C)** Insertion events are likely caused by Cas9 5’ overhang fill-in and blunt DNA EJ in all conditions. Shown is the frequency of events among the Insertion category consistent with Cas9 5’ overhang fill-in and blunt DNA EJ for distinct insertion sizes and experimental conditions. **(D)** Microhomology use (2-3 nt) is prevalent for deletion sizes -2, -7, and -13, with other deletions sizes showing no microhomology (shown is -1 to -14). Shown is the fraction for each type of microhomology for WT DMSO treated cells, which is the mean of three independent transfections. **(E)** DNA-PKcs inhibition causes an elevation in the -2 and -7 deletion sizes, which are associated with microhomology, no effect on the -13 deletion that also uses microhomology, while conversely causing a decrease in the remainder of deletion sizes that show no microhomology (shown is -1 to -14). n = 3 for each deletion size, unless a particular deletion size was not detected in at least one of the samples, Namely, §,† denote that while the mean frequencies were much lower than the WT DMSO, statistics were not feasible, because § = deletion size only found in 2 of the replicates. † = deletion size not found in any of the replicates. Statistics with unpaired t-test using Holm-Sidak correction. *P<0.05, **P<0.01, ***P<0.001, ****P<0.0001.

From this analysis, we found that loss of PAXX or MRI had similar No Indel EJ frequencies to WT cells. When WT cells were treated with 500 nM of M3814, there was a 1.8-fold decrease in No Indel EJ, and this was similar for MRI-KO cells treated with 500 nM of M3814 (2-fold decrease, Figure 2B). However, with PAXX loss (PAXX-KO and PAXX-KO/MRI-KO) M3814 caused a significantly greater decrease in No Indel EJ (4.8-fold, Figure 2B). Thus, the influence of PAXX and MRI on No Indel EJ with or without M3814 treatment was similar in this assay as for the EJ7-GFP assay. We observed a similar pattern with Insertions (Figure 2B): loss of PAXX or MRI had no effect, these events were modestly reduced with M3814 treatment in WT and MRI-KO cells, whereas in cells without PAXX, M3814 caused significant decrease in these events (2.1-fold for PAXX-KO and 1.8-fold for PAXX-KO/MRI-KO). The converse was true for deletion frequencies: M3814 caused a 1.8- and 1.5-fold increase for WT and MRI-KO cells, respectively, whereas in cells without PAXX, M3814 caused a significantly greater increase (2.5-fold for PAXX-KO and 2.6-fold for PAXX-KO/MRI-KO). Lastly, for Complex Indels, these events were much less frequent compared to the other EJ outcomes. (Figure 2B).

Since the findings with insertions were similar to No Indel EJ, we posited that the insertions are also the product of blunt DSB EJ. Specifically, we considered that the insertion mutations are primarily caused by staggered Cas9 DSBs, causing 5’ overhangs that are filled in to generate blunt DSBs prior to EJ (Supplemental Figure 2A), as shown previously (37). To test this hypothesis, we examined the sequence of insertions, and found that the vast majority of the insertions are consistent with this mechanism (Figure 2C, Supplemental Figure 2A). Notably, the +2 nucleotide insertions were the most frequent (Supplemental Figure 2B), and loss of PAXX or MRI, as well as DNA-PKcs kinase inhibition, had no effect on the frequency of the insertion size or whether the insertions were consistent with a Cas9 5’ overhang (Figure 2C, Supplemental Figure 2B). Thus, the vast majority of insertions appear to result from blunt DSB EJ, and likely explains the similar frequency patterns between insertion mutations and No Indel EJ.

We then examined the deletion EJ outcomes in more detail. Specifically, we determined the frequency of each deletion size, as well the use of microhomology at each deletion size. Beginning with WT cells treated with DMSO, we found that the most frequent deletions sizes were -2, -7, and -13, and each of these deletion sizes predominantly involved microhomology (i.e, 2 or 3 nt, Figure 2D, 2E). We also observed microhomology with some of the larger deletions, i.e., -17, -20, and -75 (Supplemental Figure 3A). The rest of the deletion sizes showed predominantly no microhomology (i.e., 0 or 1 nt, Figure 2E, Supplemental Figure 3B). Notably, M3814 treatment caused a significant increase in the frequency of deletion sizes -2, -7, -17, and -75, each of which show use of microhomology, and a converse decrease in the deletion sizes showing no microhomology (Figure 2E, Supplemental Figure 3). There was an exception to this pattern, in that the -13 deletion that also shows microhomology did not increase with M3814, and indeed showed a modest reduction (Figure 2E). Importantly, loss of PAXX and/or MRI had no substantial effects on deletion patterns with or without M3814, although for the -2 deletion, MRI loss (i.e. MRI-KO and PAXX-KO/MRI-KO) caused a decrease vs. WT M3814 (Figure 2E, Supplemental Figure 4A-C). Namely, with M3184, loss of MRI caused a reduction in the -2 nt deletion compared to WT cells treated with M3814, which involves microhomology. Altogether, these findings indicate that M3814 treatment causes a substantial increase in short deletions with microhomology and a converse reduction in deletions without microhomology. Lastly, PAXX and MRI did not show an obvious effect on these deletion patterns, however there were a few individual events (e.g., the -13 deletion) that are exceptions to this overall conclusion.

### Combining loss of DNA-PKcs with loss of PAXX causes a reduction in No Indel EJ, but otherwise has minimal effects on EJ patterns

Since loss of PAXX, but not MRI, magnified the effects of DNA-PKcs kinase inhibition on blunt DSB EJ, we then tested whether PAXX may have similar effects combined with genetic loss of DNA-PKcs. To test this, we generated a double knockout cell line (PRKDC-KO/PAXX-KO) from a previously described PRKDC-KO cell line (22), and examined EJ outcomes, using the MA-del assay (Figure 3A). For No Indel EJ, we found that loss of DNA-PKcs caused a 1.2-fold decrease, and as described above, PAXX loss did not cause a decrease (Figure 3A). However, PAXX loss in the PRKDC-KO cell line caused a 1.7-fold decrease in No Indel EJ (PRDKC-KO/PAXX-KO vs. PRKDC-KO, Figure 3A). For insertions, we found that loss of DNA-PKcs had no obvious effect (albeit a significant 1.1-fold reduction with PAXX-KO, Figure 3A), and that the insertions remained consistent with staggered Cas9 DSBs followed by blunt DSB EJ (Figure 3B, Supplemental Figure 5A). Deletion frequencies were modestly elevated in PRKDC-KO cells and only minimally changed with loss of PAXX (1.2-fold, Figure 3B). Finally, complex indels were rare in each condition. Altogether, loss of PAXX amplified the effect of genetic loss of DNA-PKcs for No Indel EJ, but to a lesser extent than the effects of PAXX combined with M3814. Furthermore, genetic loss of DNA-PKcs had modest effects on other EJ outcomes with or without PAXX, which is distinct from our findings with M3814. We suggest that the importance of PAXX for blunt EJ is greater with DNA-PKcs kinase inhibition vs. genetic loss of DNA-PKcs.

**Figure 3.**
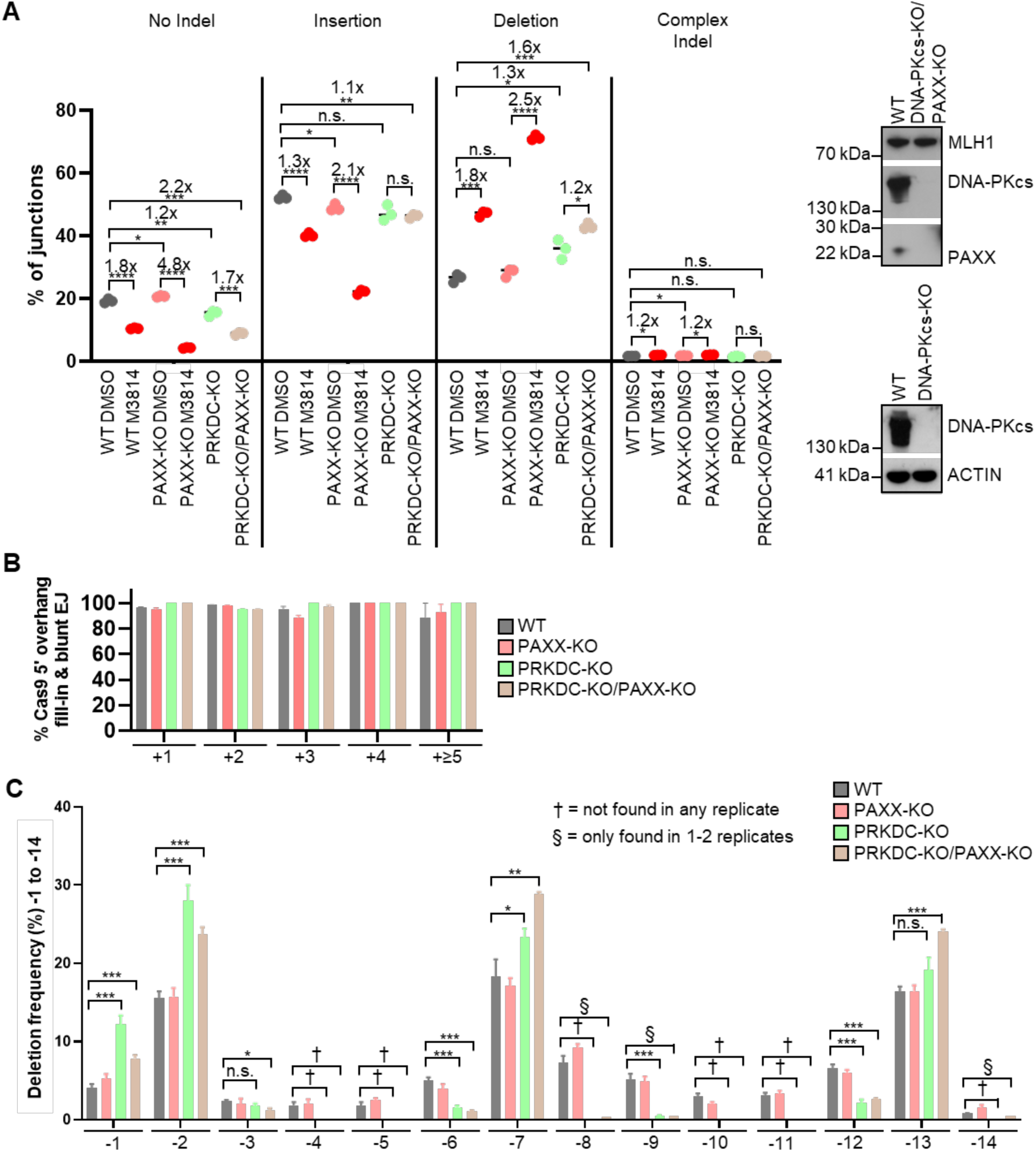
Combining loss of DNA-PKcs with loss of PAXX causes a reduction in No Indel EJ, but otherwise has minimal effects on EJ patterns. **(A)** Combined loss of DNA-PKcs and PAXX has causes a reduction in No Indel EJ and a modest increase in Deletion frequencies. Shown are junction frequencies for No Indel EJ, Insertions, Deletions, and Complex Indels for each condition. n=3 biologically independent transfections. Statistics with unpaired t-test using Holm-Sidak correction. *P<0.05, **P<0.01, P<0.001, ****P<0.0001, n.s. = not significant. WT and PAXX-KO are the same as in Figure 2B. Immunoblots show expression levels of DNA-PKcs and PAXX **(B)** Insertions are consistent with Cas9 5’ overhang fill-in and blunt EJ irrespective of DNA-PKcs and/or PAXX loss. Shown is the frequency of events among the Insertion category consistent with Cas9 5’ overhang fill-in and blunt DNA EJ for distinct insertion sizes and experimental conditions. **(C)** Loss of DNA-PKcs causes an increase in -2 and -7 deletions, which are associated with microhomology, a decrease in most deletion sizes without microhomology, except for the -1 deletion, while loss of PAXX does not cause a consistent/substantial pattern. n = 3 for each deletion size unless otherwise depicted, as in Fig 2E. §,† denote that while the mean frequencies were much lower than the WT DMSO, statistics were not feasible, because § = deletion size only found in 1-2 replicates. † = deletion size not found in any of the replicates. WT and PAXX-KO are the same as in Figure 2E. Statistics with unpaired t-test using Holm-Sidak correction. *P<0.05, **P<0.01, ***P<0.001, n.s. = not significant.

We also examined deletion sizes and found that DNA-PKcs genetic loss had similar effects as M3814, but with some minor differences. As with M3814, loss of DNA-PKcs caused an increase in the -2 and -7 deletions that are associated with microhomology, and a decrease in other deletion sizes that show no evidence of microhomology (i.e., -3 to -6, -8 to -12, and -14, Figure 3C, Supplemental Figure 5B, 5C). Loss of DNA-PKcs had no significant effect on the -13 deletion that is also associated with microhomology, which was similar to M3814. In contrast, loss of DNA-PKcs caused a significant increase in the -1 deletion that shows no microhomology, which was the opposite of M3814. Finally, combining loss of PAXX with PRKDC-KO caused a modest increase in the -7 and -13 deletions, which again are associated with microhomology (Figure 3C, Supplemental Figure 5D). These findings indicate that loss of DNA-PKcs causes elevated deletions with microhomology, and a decrease in several deletions without microhomology, which are similar to M3814, but also caused an increase in very short (1 nt) deletions without microhomology. In summary, for microhomology use during deletions, the effects of DNA-PKcs loss and kinase inhibition are similar (Figure 3C), but kinase inhibition has a greater effect on blunt DSB EJ, particularly when combined with loss of PAXX (Figure 3A).

### Combining XLF loss, or a hypomorphic allele of XLF, with loss of MRI or PAXX causes a reduction in No Indel EJ

Next, we examined the effects of PAXX and MRI on No Indel EJ when combined with disruption of XLF, by generating double KO cell lines, and using the EJ7-GFP reporter. Prior studies found that combined loss of XLF with PAXX or MRI causes a defect in V(D)J recombination, which is not observed with the single mutants (27–30). However, while XLF is dispensable for V(D)J recombination, this factor is critical for No Indel EJ of Cas9 DSBs, which we repeated here (14-fold, Figure 4A). Nonetheless, XLF-KO cells show residual No Indel EJ, which we found was further reduced with loss of either MRI (20-fold) or PAXX (242-fold), and was reversed with transient complementation of these factors (Figure 4A).

**Figure 4.**
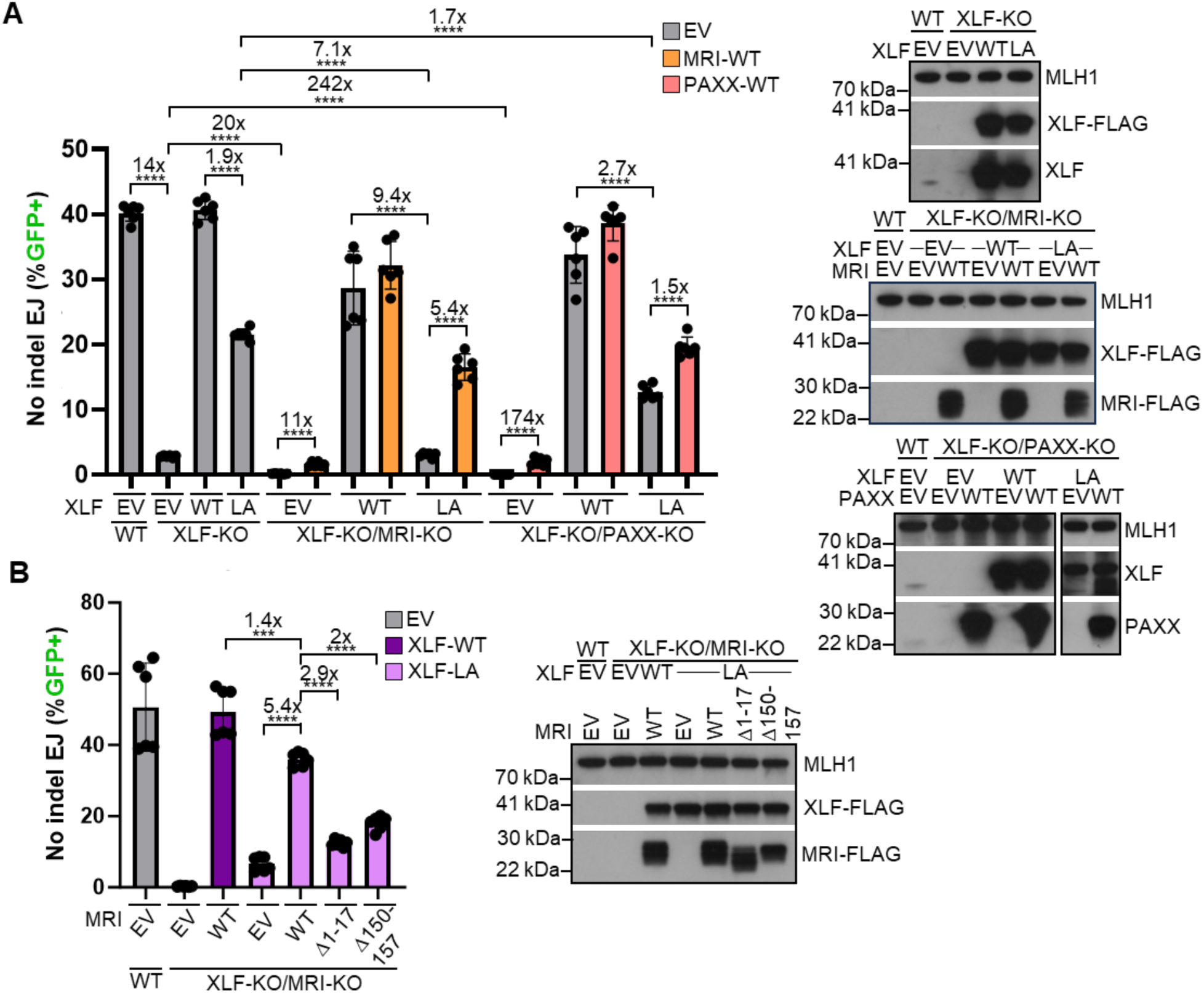
Combining XLF loss, or a hypomorphic allele of XLF, with loss of MRI or PAXX causes a reduction in No Indel EJ. **(A)** XLF loss, or a hypomorphic allele of XLF, reveal a role for MRI and PAXX to promote No Indel EJ. XLF-LA = L115A/K288A/K290A/K292A/K293A. n=6 biologically independent transfections. Statistics with unpaired t-test using Holm-Sidak correction. ****P<0.0001. Immunoblots show expression levels of XLF-FLAG, MRI-FLAG, XLF, and PAXX. **(B)** The N- and C-terminal ends of MRI are each important to promote No Indel EJ with a hypomorphic allele of XLF. n=6 biologically independent transfections. Statistics with unpaired t-test using Holm-Sidak correction. ***P<0.001, ****P<0.0001. Immunoblots show expression levels of XLF-FLAG and MRI-FLAG.

We then posited that PAXX and MRI would promote No Indel EJ not only with XLF loss, but also with a hypomorphic allele of XLF. Specifically, we used the XLF-LA (L115A/K288A/K290A/K292A/K293A) mutant, which weakens its ability to interact with XRCCC4 (L115A) and Ku80 (the C-terminal K to A mutations) (20, 38–41). The XLF-LA mutant showed a 1.9-fold decrease in No Indel EJ compared to XLF-WT (Figure 4A). This hypomorphic allele of XLF was shown previously to magnify the role of DNA-PKcs for No Indel EJ, and we considered whether this effect would also be observed with MRI and PAXX. Indeed, we found that without MRI, the XLF-LA mutant showed a 9.4-fold decrease in No Indel EJ vs. XLF-WT, which was complemented 5.4-fold with expression of MRI-WT (Figure 4A). Without PAXX, the fold-effect of the XLF-LA mutant vs. XLF-WT was also magnified (2.7-fold), and complemented 1.5-fold with expression of PAXX (Figure 4A). Thus, both MRI and PAXX are important for No Indel EJ when combined with loss of XLF and a hypomorphic allele of XLF, although the fold effect for MRI was greater than PAXX.

Since expression of MRI caused a substantial increase in No Indel EJ in cells expressing XLF-LA (in XLF-KO/MRI-KO cells), we investigated domains of MRI important for this function. Specifically, we examined the N-terminal (MRI-Δ1-17) and C-terminal (MRI Δ150-157) ends, which contain the APLF-like and XLF-like Ku binding motifs (KBMs), respectively (27). Although only the N-terminal motif has been shown to bind the Ku heterodimer (26, 27). As above, when we expressed MRI-WT with XLF-LA (in the XLF-KO/MRI-KO cells), No Indel EJ frequencies increased 5.4-fold (Figure 4B). In contrast, MRI-Δ1-17 or MRI Δ150-157 showed reduced No Indel EJ vs. MRI-WT (2.9-fold and 2-fold lower, respectively, Figure 4B). Thus, both the N-terminal and C-terminal ends of MRI are important to promote No Indel EJ in cells with a hypomorphic XLF.

### XLF loss causes a defect in EJ outcomes and is further enhanced with combined loss of MRI or PAXX

We next tested the above hypothesis using the MA-del assay to examine various EJ outcomes in XLF-KO, XLF-KO/MRI-KO, and XLF-KO/PAXX-KO HEK293 cells. From this analysis we found that loss of XLF led to a 1.9-fold decrease in No Indel EJ events (Figure 5A), whereas combined loss of XLF with MRI or PAXX substantially decreased these events (26-fold and 37-fold decrease, respectively). We found a similar pattern for insertion events, such that loss of XLF leads to a 3.7-fold decrease, but when combined with loss of MRI or PAXX led to a 17-fold or 14-fold decrease respectively. (Figure 5A). The converse was true with deletion events where XLF-KO cells caused a 2.8-fold increase in these events and was further enhanced by XLF-KO/MRI-KO and XLF-KO/PAXX-KO cells (3.5-fold increase with both, Figure 5A). Lastly, Complex Indel events were much more infrequent and loss of XLF did not significantly affect these outcomes.

**Figure 5.**
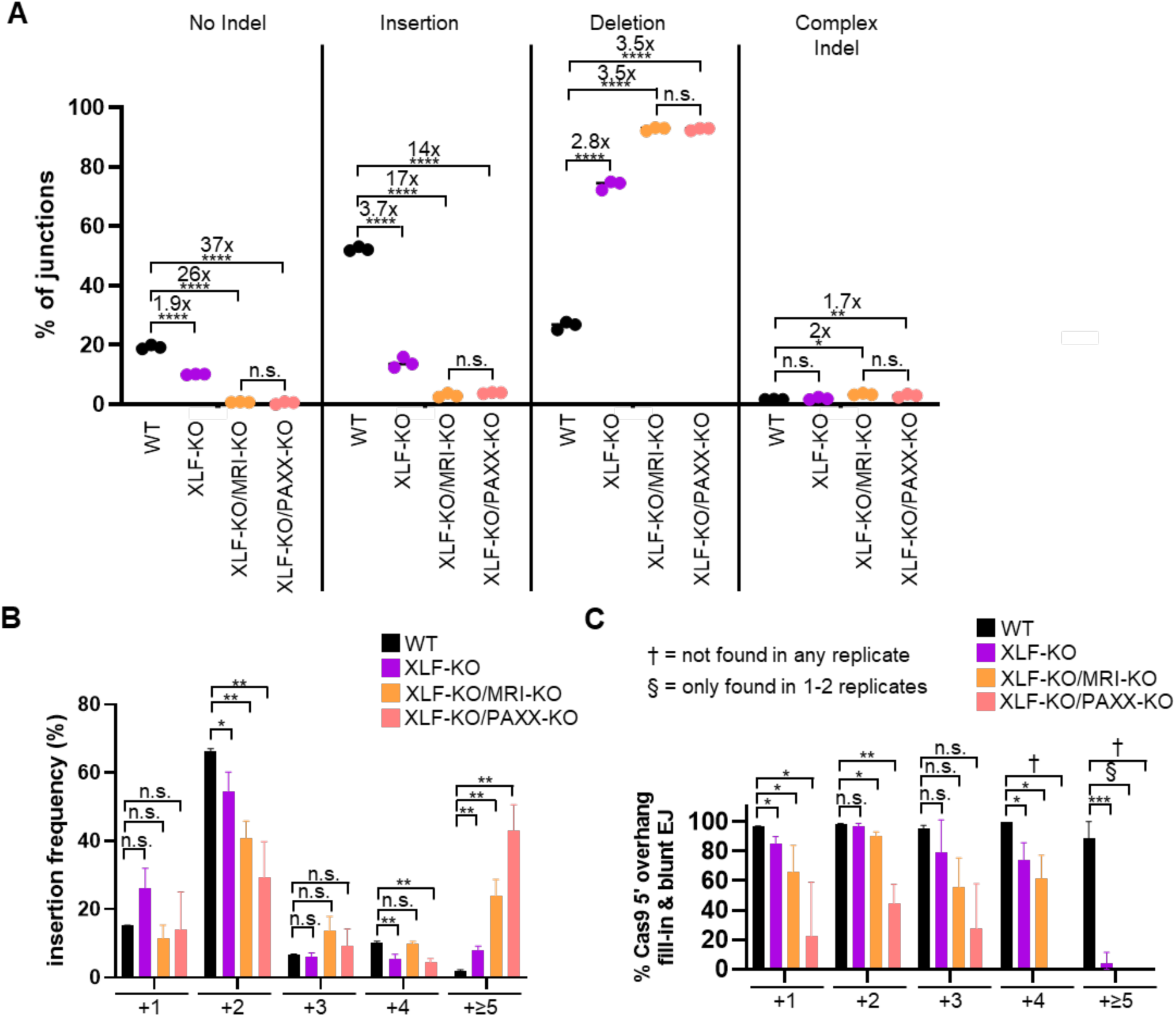
XLF loss causes a defect in EJ outcomes and is further enhanced with combined loss of MRI or PAXX. **(A)** Loss of XLF leads to a decrease in No Indel EJ and Insertions and a converse increase in Deletions, and these effects are greater when combined with loss of MRI or PAXX. Shown are junction frequencies for No Indel EJ, Insertions, Deletions, and Complex Indels for each condition. n=3 biologically independent transfections. Statistics with unpaired t-test using Holm-Sidak correction. **P<0.01, ****P<0.0001, n.s. = not significant. WT values are those seen in Figure 2B **(B)** Combined loss of XLF with MRI or PAXX causes a shift on insertion size frequencies. Statistics with unpaired t-test using Holm-Sidak correction. *P<0.05, **P<0.01, n.s. = not significant. WT values are those seen in Supplemental Figure 2B. **(C)** Combined loss of XLF with MRI or PAXX leads to a reduction in insertions consistent with Cas9 5’ overhang fill-in and blunt DNA EJ. Shown is the frequency of Insertions that are consistent with Cas9 5’ overhang fill- in and blunt DNA EJ. WT values are those seen in Figure 2C.

We further examined the insertion EJ events. Beginning with insertion size frequencies, we observed changes to the pattern among the cell lines, with the most striking difference being a significant decrease in +2 nucleotide, and a significant increase in +≥5 nt insertions for all three mutant lines, particularly for XLF-KO/MRI-KO and XLF-KO/PAXX-KO cells (Figure 5B). Notably, the +≥5 nt insertions in these mutant cell lines are also rarely consistent with Cas9 5’ overhang followed by fill-in and blunt DSB EJ (Figure 5C). Indeed, for several of the other insertion sizes, we found that loss of XLF with MRI or PAXX had significant decrease in insertions that were consistent with Cas9 5’ overhang followed by fill-in and blunt DSB EJ (Figure 5C). Thus, both the frequency of insertions, and the percentage of insertions consistent with blunt DSB EJ, were decreased with loss of these factors. Altogether, the findings with No Indel EJ and insertions indicate that loss of XLF causes a reduction in blunt DSB EJ that is magnified by combined loss of either PAXX or MRI.

### XLF loss causes a decrease in short deletions and increase in large deletions with microhomology, which is magnified by MRI loss

Finally, we examined the deletion events in more detail. We found that XLF loss caused a significant reduction in short deletion sizes without microhomology, which was similar to our findings with DNA-PKcs inhibition or loss. However, XLF loss did not significantly affect -2 and -7 nt deletions (Figure 6A), which is different to DNA-PKcs inhibition or loss, which caused a significant increase in these microhomology-associated deletion events (Figure 2E and Figure 3C). Instead, loss of XLF caused a significant increase in larger deletions such as -17, -20, and -31 nt deletions (Figure 6B), each of which involve microhomology (Figure 6C). Furthermore, we found deletion sizes in XLF deficient cells that were not present in WT cells (Figure 6B and Supplemental Figure 6A). For example, XLF deficient cells show a substantial frequency of -35 nt deletion size events, which show substantial microhomology use (Figure 6B, 6C). However, this deletion size was not present in WT cells or cells with DNA-PKcs disruption (Figure 6B, Supplemental Figures 3A, 5C).

**Figure 6.**
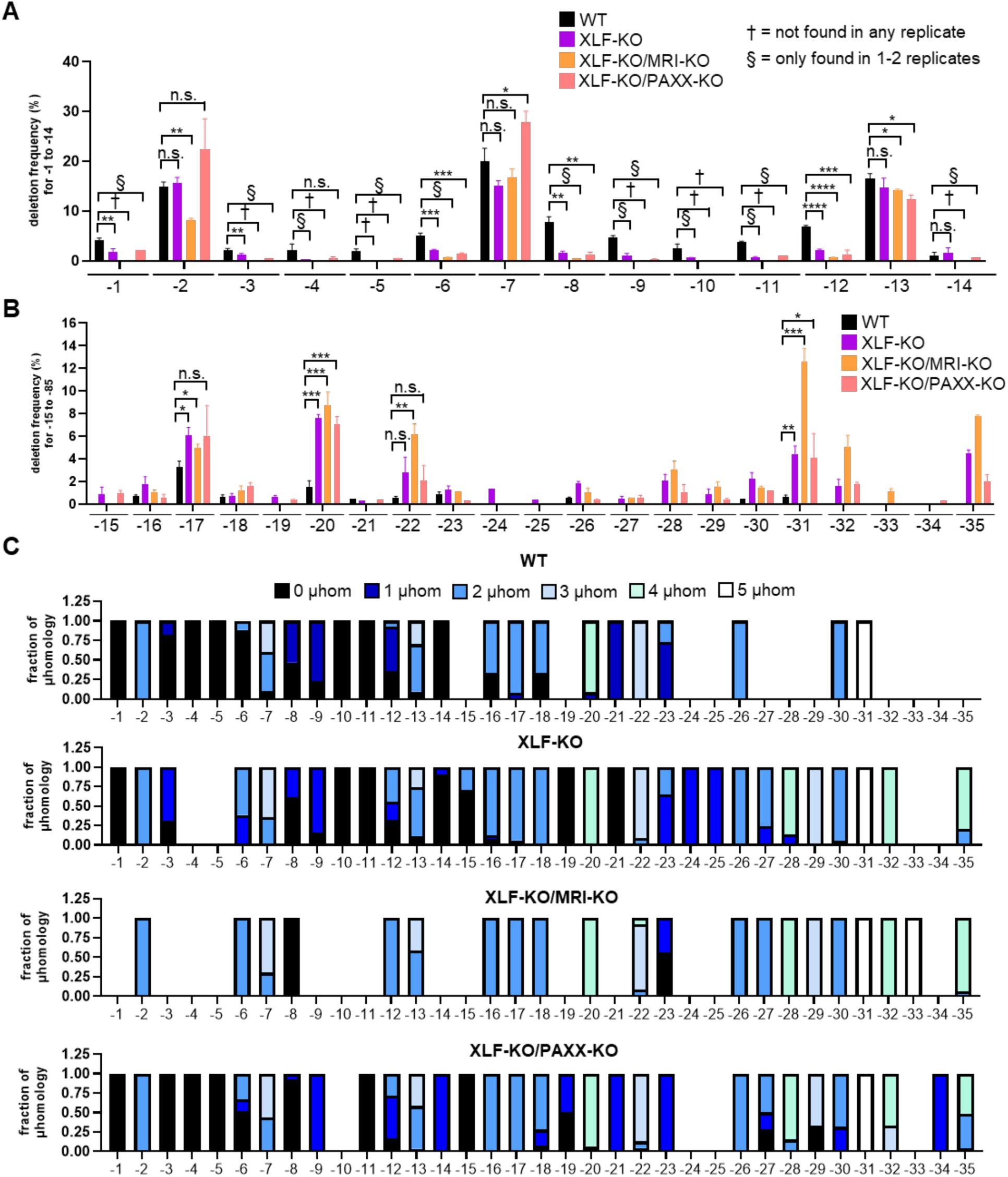
XLF loss causes a decrease in short deletions and increase in large deletions with microhomology, which is magnified by MRI loss. **(A)** Loss of XLF causes a reduction or loss in short deletions (-1 to -14) that is further enhanced by loss of MRI. n = 3 for each deletion size, unless a particular deletion size was not detected in at least one of the samples, Namely, §,† denote that while the mean frequencies were much lower than the WT, statistics were not feasible, because § = deletion size not found in all three replicates. † = deletion size not found in any of the replicates. Statistics with unpaired t-test using Holm-Sidak correction. *P<0.05, **P<0.01, ***P<0.001, ****P<0.0001, n.s. = not significant. WT values are those seen in Figure 2E. **(B)** Loss of XLF causes an increase in large deletions (-15 to -35) that is further enhanced by loss of MRI. n = 3 for each deletion size, unless a particular deletion size was not detected in at least one of the samples, Namely, §,† denote that while the mean frequencies were much lower than the WT, statistics were not feasible, because § = deletion size only found in 1-2 replicates. † = deletion size not found in any of the replicates. Statistics with unpaired t-test using Holm-Sidak correction. *P<0.05, **P<0.01, ***P<0.001, n.s. = not significant. WT values are those seen in Supplemental Figure 3C. **(C)** Fraction of microhomology associated with each deletion size (-1 to -35) in WT, XLF-KO, XLF-KO/MRI-KO, and XLF-KO/PAXX-KO cells. Shown is the average fraction from for each microhomology. WT values are those seen in Figure 2D and Supplemental Figure 3A.

With the double mutants, combined loss of XLF and MRI caused a marked increase in some large deletions with microhomology vs. XLF-KO and WT (i.e., -22, -31, -35. Figure 6B, 6C). Finally, combining loss of XLF with loss of PAXX did not show a marked difference vs. XLF loss alone. Overall, we found that loss of XLF leads to a different profile of deletion sizes vs. DNA-PKcs disruption, and several deletion events caused by XLF loss were further elevated when combined loss of MRI (25).

## DISCUSSION

We sought to understand the role of PAXX and MRI during chromosomal EJ. To do so, we used Cas9 based tools measuring repair between two tandem DSBs: the EJ7-GFP reporter for No Indel EJ, and the MA-del assay to measure diverse EJ outcomes. We found that loss of PAXX and MRI led to minor defects during EJ, which is consistent with other studies (28, 29, 42). Thus, we tested possible redundancies with other factors, finding that inhibition and loss of DNA-PKcs revealed a role for PAXX for No Indel EJ, and loss and a hypomorphic allele of XLF revealed roles for both PAXX and MRI for No Indel EJ. Given the roles of DNA-PKcs and XLF in DSB end synapsis (13, 14), these findings indicate that PAXX and MRI may also contribute to this process. We also identified several distinctions, particularly regarding deletion mutation patterns, which we suggest provides an additional mechanistic insight of PAXX and MRI during chromosomal EJ.

For one, while the overall pattern of DNA-PKcs loss vs. kinase inhibition were similar with loss of PAXX, DNA-PKcs loss has a milder defect compared to DNA-PKcs kinase inhibition. These findings are consistent with the kinase-dead alleles of DNA-PKcs causing greater genome instability phenotypes vs. loss, as is also found with ATM and ATR kinases (43). DNA-PKcs kinase inhibition blocks autophosphorylation of DNA-PKcs, which is critical step for DNA-PKcs dissociation from DNA ends (13, 33–35). Such removal of DNA-PKcs in the LR complex is important for the transition to the SR complex that responsible for DNA end ligation (13, 14). Thus, DNA-PKcs kinase inhibition likely disrupts this LR to SR transition (13, 14, 33–35). In contrast, with loss of DNA-PKcs, the SR complex could form *de novo* to support efficient blunt DSB ligation (13). Since loss of PAXX had the greatest effect on blunt DSB EJ when combined with DNA-PKcs kinase inhibition, and since PAXX appears to support DSB end synapsis within the LR complex with DNA-PKcs (25, 44), we suggest that PAXX-mediated end synapsis supports the transition between the LR and SR complex. Consistent with this notion, we found that the PAXX KBM, which likely mediates such synapsis via interaction with Ku70 (25, 42, 44), is important for this function. The role of PAXX in such a transition could be via supporting end synapsis during displacement of DNA-PKcs prior to engagement with XLF-mediated synapsis. Accordingly, it is possible that PAXX also supports end synapsis after DNA-PKcs displacement, and hence is a part of the SR complex. This model is supported by our findings that PAXX promotes No Indel EJ in cells with DNA-PKcs loss. Furthermore, PAXX promotes No Indel EJ when combined with XLF loss and the XLF-LA mutant, which also may suggest a role in the SR complex, in which end synapsis appears to be primarily mediated by the XLF homodimer (24, 44). However, it is unclear from current structural studies whether PAXX can form complexes without DNA-PKcs (i.e., whether PAXX can be found in the SR complex). Thus, structural studies of SR complexes with PAXX would likely provide further insight into its function.

We note that a prior study in mouse embryonic stem cells (mESCs) found that loss of PAXX caused a modest (1.5-fold), but significant reduction in No Indel EJ without disruption of DNA-PKcs, which is different from our findings with here with HEK293 cells (20). Similarly, the XLF-LA mutant shows a greater defect in mESCs vs. HEK293 (20, 22). While the mechanism for these distinctions are unclear, one cause may be the substantially lower levels of DNA-PKcs in rodent cells vs. human, which have been estimated at 50-fold (45).

To provide a contrast to PAXX, we also examined the role of the Ku-binding factor MRI, finding that disruption of XLF, but not DNA-PKcs, revealed a role for MRI for No Indel EJ. Namely, combined loss of XLF and MRI (XLF-KO/MRI-KO) led to ablation of No Indel EJ. In addition, when hypomorphic allele of XLF is combined with loss of MRI, this led to a dramatic decrease in No Indel EJ. These findings are consistent with the defects in V(D)J recombination caused by combined loss of MRI and XLF (27–29). In contrast, MRI, also called CYREN, was shown to inhibit C-NHEJ events for DSB ends with overhangs (e.g., telomere fusions in S/G2), which indicates that the structure of DSB ends may influence the relative effect of MRI on C-NHEJ (46). MRI is very small peptide and is mainly composed of disordered regions (27, 47), and also contains two KBMs, one of which is an XLF-like KBM and the other an APLF-like KBM (26). Notably, we found mutants that disrupt these KBM regions of MRI were unable to promote No Indel EJ compared to MRI-WT. Recent studies have implicated the XLF KBM in forming molecular condensates that facilitate C-NHEJ (48, 49). Thus, it is possible that MRI may plays a similar role in stabilizing LR or SR complexes. Thus, future structural and biophysical analysis of MRI, particularly with XLF hypomorphic proteins, may provide insight into its role in blunt DSB EJ.

Apart from blunt DSB EJ, we also examined the patterns of deletion mutations, and found substantial differences between disruption of DNA-PKcs vs. XLF. DNA-PKcs kinase inhibition and loss both caused an increase in short deletion mutations with microhomology. Although as an exception to this pattern, two larger deletion sizes were also increased (i.e., -17 and -75). Thus, in addition to its role in promoting end synapsis in the LR complex, DNA-PKcs also appears to regulate processing of Cas9 DSB ends, which is consistent with its established role in promoting processing of hairpin coding ends during V(D)J recombination (4, 50–52). In contrast, we found that loss of XLF caused an increase in larger deletion mutations that were associated with microhomology, and a decrease in short deletions with microhomology. Apart from this distinction, both disruption of DNA-PKcs and XLF had the same effect on causing a decrease in deletions without microhomology. In summary, while DNA-PKcs and XLF both suppress deletions with microhomology, the deletion size is different, with DNA-PKcs being particularly important to suppress short deletions. We speculate that loss of XLF likely disrupts both the LR and SR complex, causing a bias towards longer DNA end resection and in turn lead to larger deletion mutations. In summary, these results suggest that distinct aspects of the C-NHEJ complex affect different end processing events, which then affects EJ outcomes.

Finally, we examined the effects of PAXX and MRI on deletion mutation spectra. With PAXX, we did not observe a substantial effect on deletion mutation patterns when combined with XLF disruption. Furthermore, although DNA-PKcs disruption revealed a role for PAXX on blunt DSB EJ, we did not observe a role of PAXX on deletion patterns with or without DNA-PKcs disruption. Thus, we suggest that PAXX has a significant role on DNA end synapsis during EJ rather than regulation of end processing *per se*. In contrast, loss of MRI magnified multiple effects of XLF loss: both a defect in blunt DSB EJ, as well as elevated large deletions with microhomology. As above, we speculate that MRI may play a stabilizing role in C-NHEJ complexes that becomes apparent with XLF loss. Altogether, we have shown that PAXX and MRI have distinct roles in relation to the C-NHEJ machinery that has substantial effects on the outcomes of repair.

## METHODS

### Plasmids and Cell Lines

The px330 plasmid was used for sgRNA/Cas9 expression (Addgene 42230, generously deposited by Dr. Feng Zhang). Supplemental Table 1 shows the sequences for sgRNAs used in this study. The pCAGGS-3xFLAG-XLF, and pgk-puro plasmids were previously described (22). Expression vectors for 3xFLAG-MRI and 3xFLAG-PAXX were generated from gBLOCKs (Integrated DNA Technologies) and cloned into a pCAGGS-BSKX (22), which is the empty vector (EV) control.

The parental cell line HEK293 Flp-In T-REx EJ7-GFP was previously described(22). The XLF-KO and PRKDC-KO cell lines were also previously described(22). PAXX-KO, MRI-KO, PAXX-KO/MRI-KO, PRKDC-KO/PAXX-KO cell lines were generated using Cas9/sgRNAs cloned into px330. PAXX-KO cells were used to generate PAXX-KO/MRI-KO and PRKDC-KO cells were used to generate PRKDC-KO/PAXX-KO. The parental cell line was co-transfected with the Cas9/sgRNA plasmids and a pgk-puro plasmid, transfected cells were enriched using transient puromycin treatment, followed by plating at low density to isolate and screen individual clones for gene disruption. Both PAXX and MRI disruption used two tandem sgRNAs, such that primary screening involved PCR to identify clones with deletion mutations, which were subsequently screened by immunoblotting for PAXX, and quantitative RT-PCR for MRI (oligonucleotides shown in Supplemental Table 1).

### DSB Reporter Assays

For the EJ7-GFP reporter assay, HEK293 cells were seeded at 0.5×10^5^ cells/well in a 24-well plate, the following day, cells were transfected with 200 ng of 7a and 7b (sgRNA/Cas9 plasmids), along with 50 ng plasmid for expressing MRI, PAXX, XLF, or EV control. Transfection efficiency had parallel transfections with 200 ng of GFP expression plasmid (pCAGGS-NZE-GFP) and 200 ng of EV with respective amounts of MRI, PAXX, PAXX, or EV. Transfections used 1.8 μL Lipofectamine 2000 (Thermofisher) and 0.5 mL of antibiotic free media per well. Cells were incubated for 4 hours with the transfection, washed, and replaced with complete media containing M3814 (i.e., Nedisertib, Selleckchem #S8586) and/or vehicle (Dimethyl Sulfoxide, DMSO), with all wells having the same total amount of DMSO in each experiment. Cells were analyzed using flow cytometry 3 days after transfection (ACEA Quanteon, Agilent NovoExpresss Version 1.5.0).

For the MA-del analysis, the transfection conditions were 800 ng each of MTAP and CDK2NB-AS1 Cas9/sgRNA plasmids, 400 ng pgk-puro plasmid, and 7.2 µL Lipofectamine 2000 in 2 mL antibiotic free media on a 6 well dish. M3814 treatment was performed as for the EJ7-GFP assay. The day after transfection, cells were treated with 2 µg/mL puromycin (maintaining M3814 or DMSO treatment) for 2 days to enrich for transfected cells, and then expanded without treatments for 3 days. Genomic DNA was isolated as described(53). PCR amplification (Platinum HiFi Supermix, Thermo Fisher) of the MTAP-CDK2NB1 rearrangement used the primers MAfusion1UP and MAfusion1DN. The amplicons were subjected to deep sequencing using the Amplicon-EZ service (Azenta) and their SNP/INDEL detection pipeline, in which reads were aligned to the reference sequence (i.e., predicted No Indel EJ junction sequence based on the sgRNA/Cas9 cut sites). From this analysis, EJ outcomes were categorized as No Indel EJ, Insertions/Complex Indels, or Deletions, and the total reads in each category were used to assess their frequency. Analysis of the insertion and deletion sequences was performed on all read sequences representing at least 0.1% of the combination of insertions and complex indels, and at least 0.2% of the deletion reads, respectively. Some deletion events showed nucleotides aligned within the deletion, which were most often consistent with staggered Cas9 cleavage, and hence were used to assign breakpoints for deletion size and microhomology, but if not (found only in the PAXX-KO/XLF-KO cell line), breakpoints could not be assigned, and these events could not be used for such analysis. For each condition, deep sequencing was performed on 3 independent transfections, and frequencies represent the mean ± SD.

### Immunoblotting and quantitative reverse transcription PCR (qRT-PCR)

For immunoblot analysis, cells were transfected as described in the reporter analysis, however the sgRNA/Cas9 plasmid was replaced with the EV plasmid (pCAGGS-BSKX) and were scaled 4-fold for a 6-well plate. Transfected cells were then lysed with ELB (250 mM NaCl, 5 mM EDTA, 50 mM Hepes, 0.1% (v/v) Ipegal, and Roche protease inhibitor) with sonication (Qsonica, Q800R). For the DNAPKcs-S2056p analysis, cells were pre-treated with M3814 or vehicle (DMSO) for 3 hr, followed by a 10 Gy IR (Gammacell 3000) treatment and allowed to recover for 1 hour. Protein was extracted with ELB buffer described above plus PhosSTOP (Roche) and 50 μM sodium fluoride. Blots were probed with antibodies for PAXX (Abcam ab126353), FLAG-HRP (Sigma A8592), XLF (Bethyl A300-730A), DNA-PKcs (Invitrogen MA5-13238), DNAPKcs-S2056p (Abcam ab124918), MLH1 (Abcam ab92312), ACTIN (Sigma A2066), HRP goat anti-mouse (Abcam ab205719), and HRP goat anti-rabbit (Abcam ab205718). ECL reagent (Amersham Biosciences) was used to develop HRP signals.

For qRT-PCR analysis, RNA was extracted using the Qiagen RNAeasy kit (Cat# 74104), reverse transcribed with MMLV-RT (Promega), and cDNA was amplified with iTaq Universal SYBR Green (Biorad, 1725120) along with primers for MRI or ACTIN (Supplemental Table S1). Quantification was on the Biorad CRX Connect Real-Time PCR Detection System (#1855201). The relative levels of mRNA were determined using the cycle threshold (Ct) value for MRI and then subtracted by the average Ct ACTIN control value. This value was then subtracted from the MRI-KO Ct value to calculate the ΔCt from WT cells and was used to calculate the 2−ΔΔCt value.

## ACKNOWLEDGEMENTS

This study was funded in part by the National Cancer Institute of the National Institutes of Health: R01CA256989, R01CA240392 (J.M.S.); P30CA33572 (City of Hope Core Facilities); F99CA284248 (M.C.A.); Roberts Summer Academy (R.C.).

## DATA AVAILABILITY STATEMENT

The datasets in the study are included in the study and are available from the corresponding author on reasonable request.

