## Supplementary Material for "Distinct functions of PAXX and MRI during chromosomal end joining"

**Supplemental Table S1. Oligonucleotide list.**

| **Name** | **Purpose** | **Sequence (5' → 3', all sgRNA sequences have the initial G nucleotide, regardless of whether it is part of the targeted sequence)** |
|---|---|---|
| 7a | sgRNA | GACCACCCTGACCTACGGCTA |
| 7b | sgRNA | GGCTGAAGCACTGCACGAAT |
| MTAP | sgRNA | ggccatacagaccatggtaa |
| CDK2NB-AS1 | sgRNA | tcgtcactacgctgatggca |
| PAXXsg1 | sgRNA | CACCGcggcggcgacagcggatcca |
| PAXXsg2 | sgRNA | CACCgCAGGCTGTCCGGCGTGAAGC |
| MRIsg1 | sgRNA | CACCGctcgtcacacccggaattac |
| MRIsg2 | sgRNA | CACCGtgagatagttgatgttgctc |
| MTAP | primer | aagtgtgatgggcaagaagg |
| CDK2NB-AS1 | primer | aggattctgcacttggatgg |
| *PCR genotyping primers* | | |
| PAXX PCRup | primer | GGCTCCGGAAGTCGTTCT |
| PAXX PCRdn | primer | CAGAGGGTGTCAGAGCCTTC |
| MRI PCRup | primer | CAGAGGCTGCAGGAGGTATC |
| MRI PCRdn | primer | GGGCGTCTCTCTTACTGACG |
| *qPCR analysis primers for MRI-KO mRNA* | | |
| MRIup | primer | GTCTCCTCTGACTTCAACAGC |
| MRIdn | primer | ACCACCCTGTTGCTGTAGCCAA |
| GAPDHup | primer | GTCTCCTCTGACTTCAACAGC |
| GAPDHdn | primer | ACCACCCTGTTGCTGTAGCCAA |

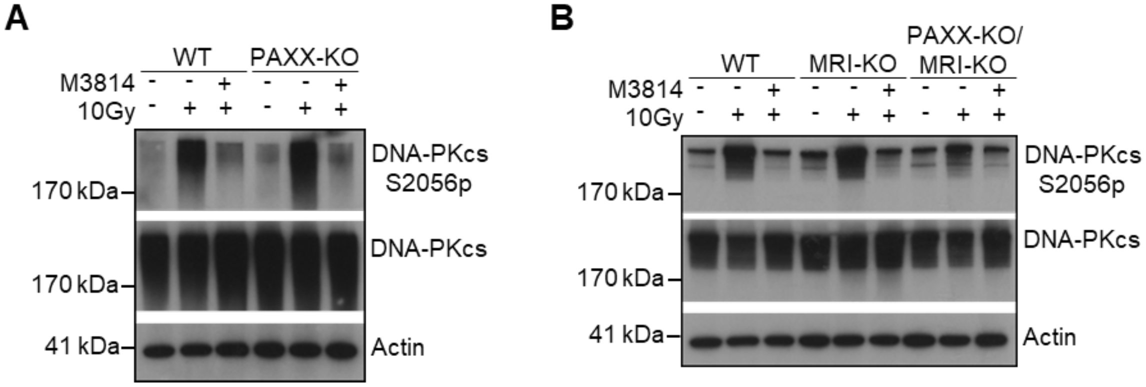


**Supplemental Figure 1. Effects of M3814 in PAXX-KO, MRI-KO, and PAXX-KO/MRI-KO HEK293 cells on DNA-PKcs pS2056. (A**) Effects of M3814 on phosphorylation of DNA-PKcs S2056 in WT and PAXX-KO cells treated with 500 nM of M3814 and 10 Gy ionizing radiation. Shown are immunoblot signals from an experiment performed for DNAPKcs-S2056p, DNA-PKcs, and Actin. **(B)** Effects of M3814 on phosphorylation of DNA-PKcs p2056 in WT, MRI-KO, and PAXX-KO/MRI-KO cells. Same treatment as above.

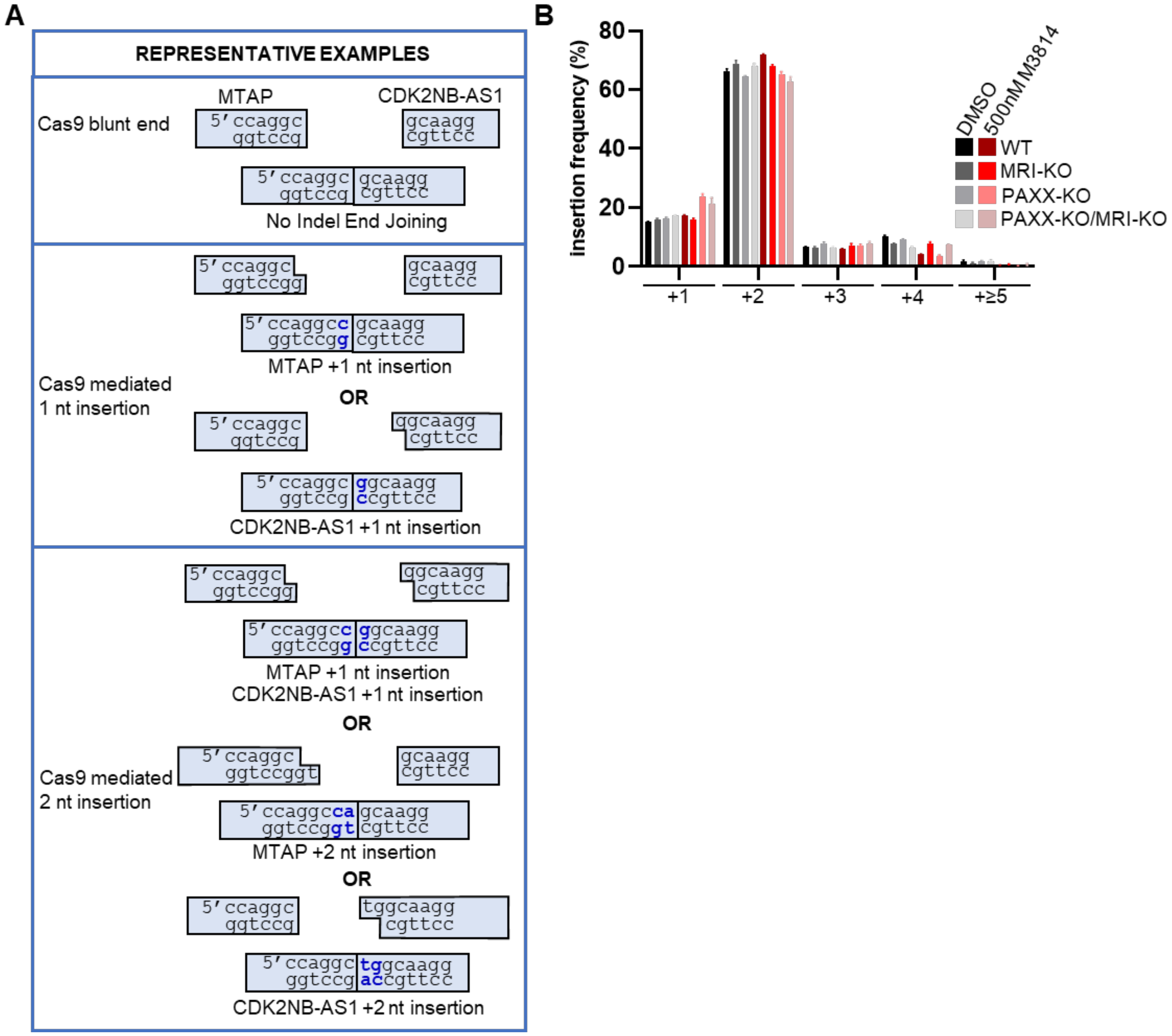


**Supplemental Figure 2. MTAP-CDK2NB-AS1 predicted insertions consistent with Cas9 staggered DSBs causing 5' overhangs and insertion size frequencies in KO HEK293 cells. (A)** Shown are the representative examples of EJ outcomes from Cas9 blunt DSB (No Indel EJ), a staggered 1 nt 5' overhang DSB at the MTAP (+1 C nucleotide insertion) or CDK2Nb-AS1 (+1 G nucleotide insertion) junction, and a staggered 2 nt 5' overhang DSB at the MTAP (+2 CA nucleotide insertion) or CDK2Nb-AS1 (+2 TG nucleotide insertion). **(B)** Insertion size frequencies in WT, MRI-KO, PAXX-KO, and PAXX-KO/MRI-KO cells treated with or without M3814. n=3 biological replicates.

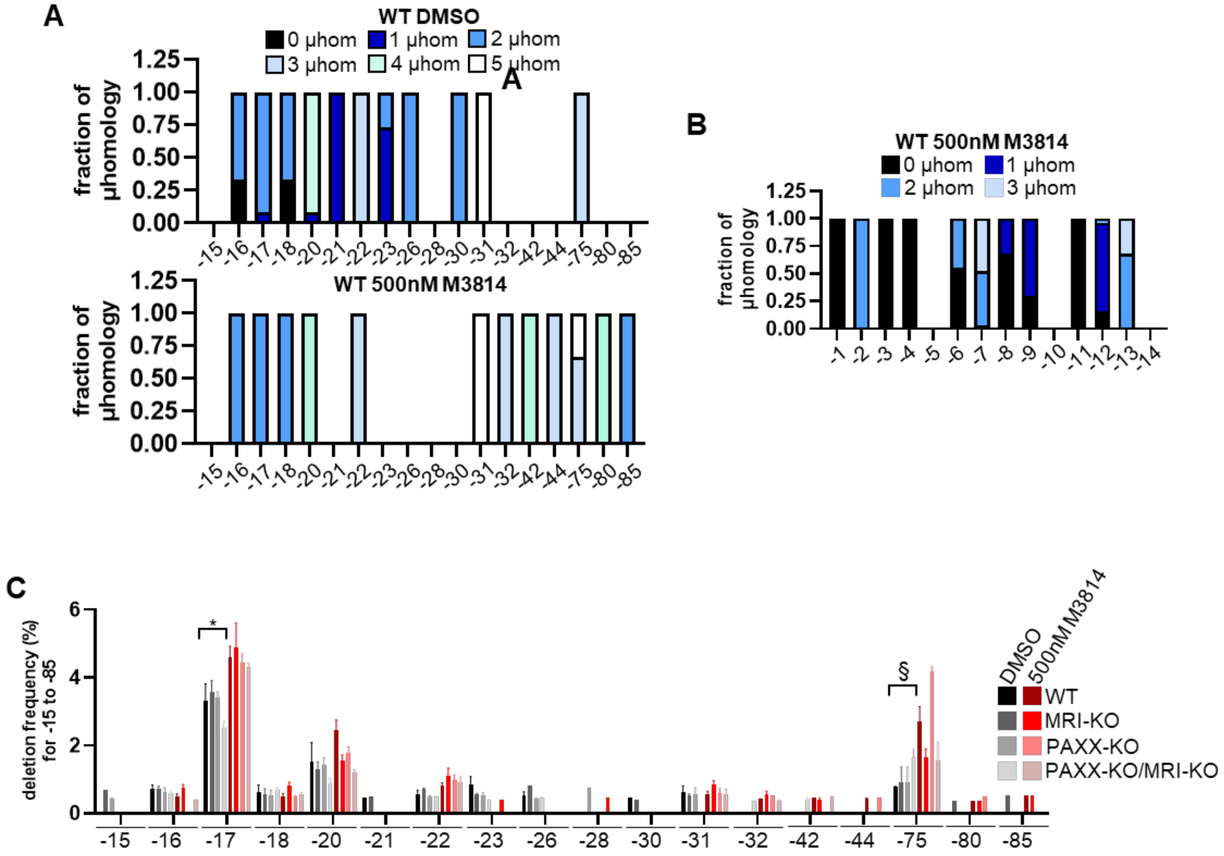


**Supplemental Figure 3. Microhomology fractions and deletion size frequencies KO HEK293 cells. (A)** Fraction of microhomology associated with each deletion size (-15 to -85) in WT cells treated with DMSO or M3814. Shown is the average fraction from for each microhomology. **(B)** Fraction of microhomology associated with each deletion size (-1 to -14) in WT 500 nM M3814 treated cells. Shown is the average fraction from for each microhomology **(C)** Deletion size frequencies (-15 to -85) in WT, MRI-KO, PAXX-KO, PAXX-KO/MRI-KO cells treated with DMSO or M3814. n = 3 for each deletion size unless otherwise depicted. § = that deletion size only found in 2 replicates for the WT DMSO cell line. Statistics with unpaired t-test using Holm-Sidak correction. *P<0.05.

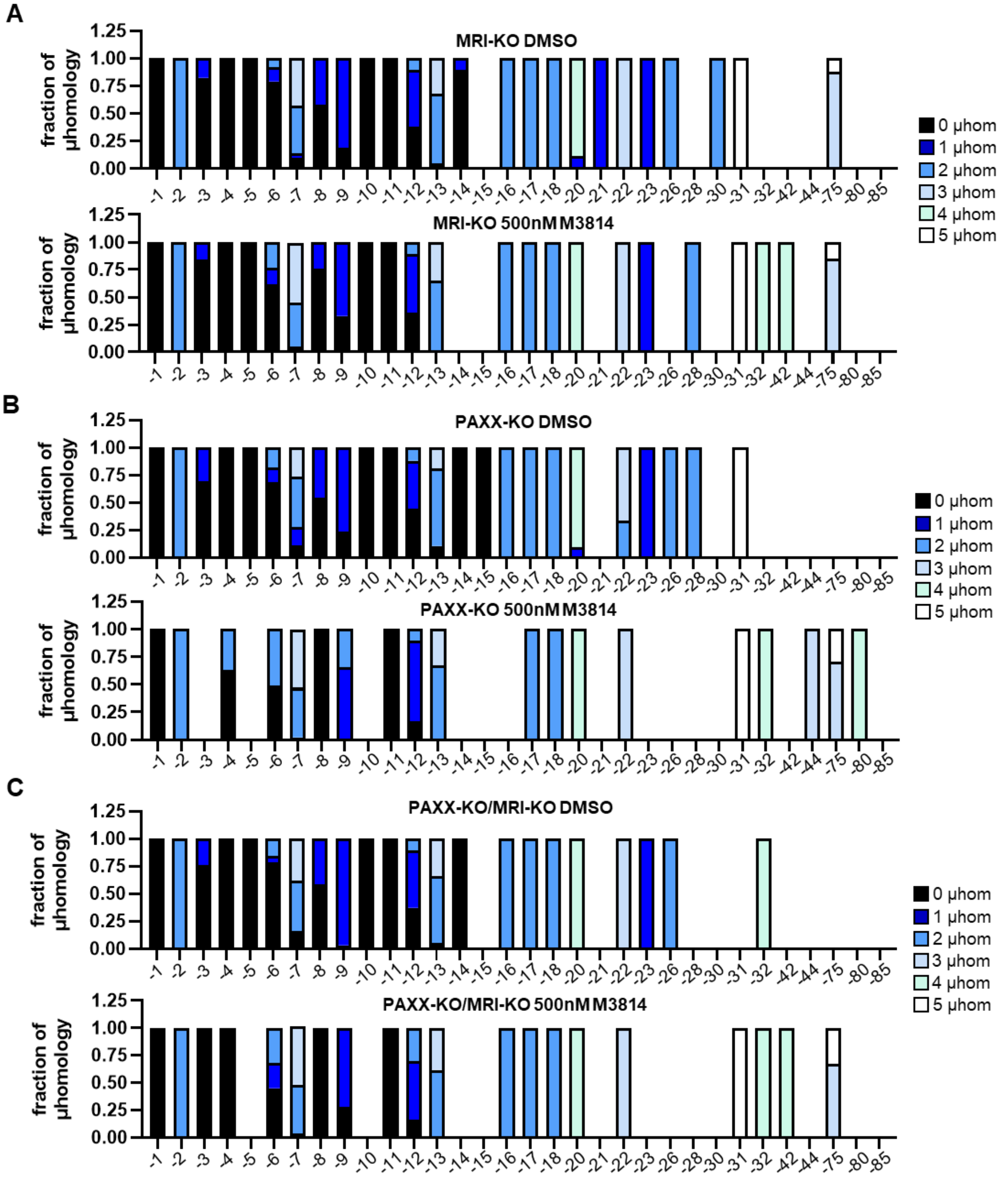


**Supplemental Figure 4. MTAP-CDK2NB-AS1 microhomology fractions MRI-KO, PAXX-KO, and PAXX-KO/MRI-KO cells. (A)** Fraction of microhomology associated with each deletion size (-1 to -85) in MRI-KO cells treated with DMSO or M3814. Shown is the average fraction from for each microhomology. **(B)** Fraction of microhomology associated with each deletion size (-1 to -85) in PAXX-KO cells treated with DMSO or M3814. Shown is the average fraction from for each microhomology. **(C)** Fraction of microhomology associated with each deletion size (-1 to -85) in PAXX-KO/MRI-KO cells treated with DMSO or M3814. Shown is the average fraction from for each microhomology.

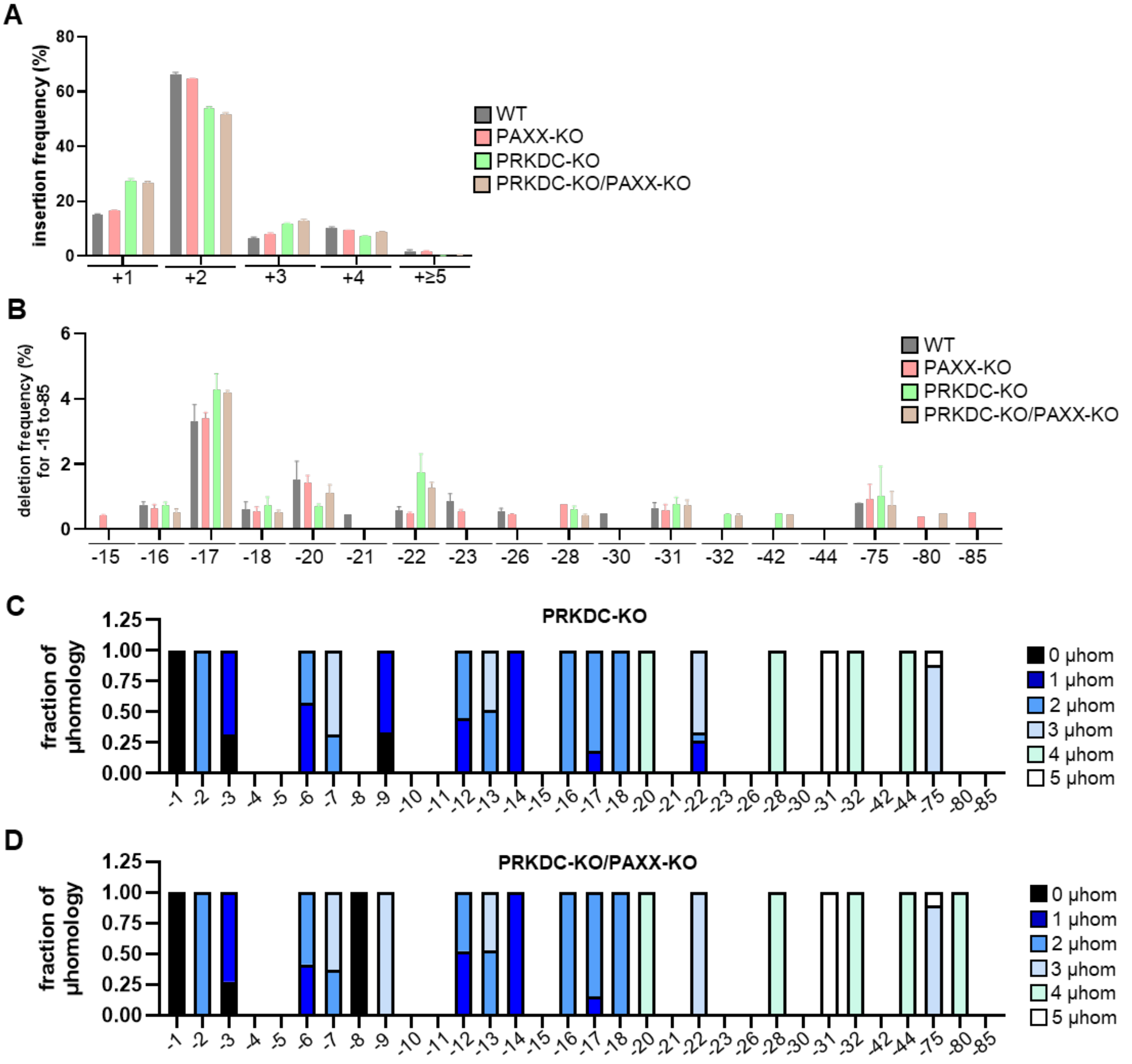


**Supplemental Figure 5. Insertions size frequencies, deletion size frequencies, and microhomology fractions in WT, PAXX-KO, PRKDC-KO, and PRKDC-KO/PAXX-KO cells.** **(A)** Insertion size frequencies in WT, PAXX-KO, PRKDC-KO, and PRKDC-KO/PAXX-KO cells. n=3 independent transfections. WT and PAXX-KO are the same values seen in Supplemental Figure 2A **(B)** Deletion size frequencies (-15 to -85) in WT, PAXX-KO, PRKDC-KO, and PRKDC-KO/PAXX-KO cells. WT and PAXX-KO are the same values seen in

Supplemental Figure 3C. **(C)** Fraction of microhomology associated with each deletion size (-1 to -85) in PRKDC-KO cells. Shown is the average fraction from for each microhomology. **(D)** Fraction of microhomology associated with each deletion size (-1 to -85) in PRKDC-KO/PAXX-KO cells. Shown is the average fraction from for each microhomology.

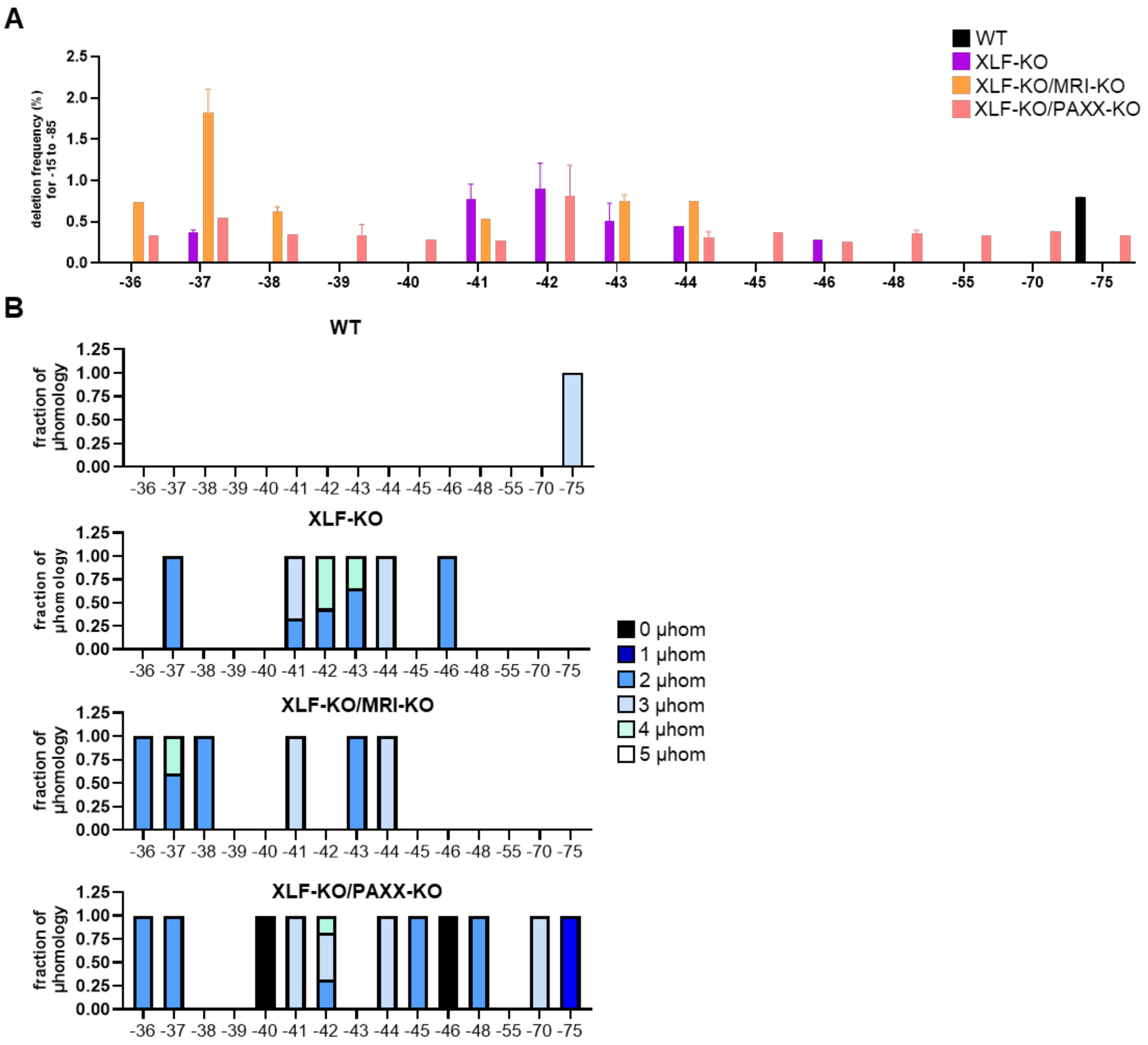


**Supplemental Figure 6. Deletion size frequencies and microhomology fractions in WT, XLF-KO, XLF-KO/MRI-KO, and XLf-KO/PAXX-KO, PRKDC-KO of -36 to -75. (A)** Deletion size frequencies (-36 to -75) in WT, XLF-KO, XLF-KO/MRI-KO, XLF-KO/PAXX-KO. WT values are those seen in Supplemental Figure 3C. **(B)** Fraction of microhomology associated with each deletion size (-36 to -75) in WT, XLF-KO, XLF-KO/MRI-KO, XLF-KO/PAXX-KO cells. Shown is the average fraction from for each microhomology. WT values are those seen in Supplemental Figure 3A.
